# Fmrp regulates oligodendrocyte lineage cell specification and differentiation

**DOI:** 10.1101/2021.03.16.435661

**Authors:** Caleb A. Doll, Kayt Scott, Bruce Appel

**Affiliations:** Department of Pediatrics, Section of Developmental Biology, University of Colorado School of Medicine, Children’s Hospital Colorado, Aurora, CO 80045, USA

## Abstract

Neurodevelopment requires the precise integration of a wide variety of neuronal and glial cell types. During early embryonic development, motor neurons and then oligodendrocyte precursor cells (OPCs) are specified from neural progenitors residing in the periventricular pMN progenitor domain of the spinal cord. Following gliogenesis, OPCs can differentiate as oligodendrocytes (OLs) – the myelinating glial cells of the central nervous system - or remain as OPCs. To generate unique cell types capable of highly divergent functions, these specification and differentiation events require specialized gene expression programs. RNA binding proteins (RBPs) regulate mRNA localization and translation in the developing nervous system and are linked to many neurodevelopmental disorders. One example is Fragile X syndrome (FXS), caused by the loss of the RBP fragile X mental retardation protein (FMRP). Importantly, infants with FXS have reduced white matter and we previously showed that zebrafish Fmrp is autonomously required in OLs to promote myelin sheath growth. We now find that Fmrp regulates cell specification in pMN progenitor cells and subsequently promotes differentiation of OPCs, such that *fmr1* mutant zebrafish embryos generate excess OPCs and fewer differentiating OLs in the developing spinal cord. Although the early patterning of spinal progenitor domains appears largely normal in *fmr1* mutants during early embryogenesis, Shh signaling is greatly diminished. Taken together, these results suggest cell stage-specific requirements for Fmrp in the specification and differentiation of oligodendrocyte lineage cells.

## Introduction

RNA binding proteins, including fragile X mental retardation protein (FMRP), play essential roles in a wide variety of neurodevelopmental processes and are linked to autism spectrum disorders (ASDs; Verkerk et al., 1991; Lee et al., 2016; Barone et al., 2017). Fragile X syndrome (FXS) is caused by a trinucleotide expansion in the 5’ UTR of the *FMR1* gene, leading to loss of FMRP expression (Verkerk et al., 1991). The RBP binds an array of mRNAs associated with ASDs (Darnell et al., 2011; Ascano et al., 2012) and can regulate the stability (Zalfa et al., 2007), transport and translation (Pilaz et al., 2016) of bound mRNAs. Though the bulk of FXS studies have centered on neuronal dysfunction (Pan et al., 2004; Dictenberg et al., 2008; Doll and Broadie, 2015; Doll et al., 2017), oligodendrocytes are also implicated in the disease state, as white matter is reduced in infants with FXS (Swanson et al., 2018), and we recently showed that human FMRP can autonomously rescue the diminished myelin sheath growth seen in zebrafish *fmr1* mutants (Doll et al., 2019). As myelin plays essential roles in neural circuit maturation and function (Mckenzie et al., 2014; Pan et al., 2020; Wang et al., 2020), this underscores the importance of FMRP in glial cell function.

Our current work highlights upstream roles for zebrafish Fmrp in the formation of oligodendrocyte precursor cells (OPCs) and subsequent differentiation into oligodendrocytes (OLs), which could contribute to the diminished myelination seen in *fmr1* mutants. We focus on the ventral spinal cord, where progenitor cells in the pMN domain sequentially generate spinal motor neurons and then OPCs. Importantly, Sonic hedgehog (Shh) signaling regulates both the transition from neurogenesis to gliogenesis in the pMN and the subsequent differentiation of OPCs to myelinating OLs (Dessaud et al., 2007; Ravanelli et al., 2018; Scott et al., 2020). However, not all OPCs differentiate into myelinating oligodendrocytes and these precursors represent a distinct, highly dynamic, and physiologically active cell type that persist into adulthood (Bergles et al., 2000; Dawson et al., 2003a; Tanaka et al., 2009).

What regulates the choice between myelin-fated OPCs vs persistent OPCs? As these cell types display widely divergent gene expression profiles (Perlman et al., 2020), RNA binding proteins must provide essential post transcriptional regulation of nascent mRNA in the oligodendrocyte lineage. Importantly, *Fmr1* deletion leads to a fewer neurons and a surplus of neural progenitor cells (NPCs; Edens et al., 2019), thereby implicating the RBP in neuronal differentiation. Moreover, FXS models show disproportionate ratios of cell types throughout the nervous system, including excess glutamatergic neurons and astrocytes (Tervonen et al., 2009; Luo et al., 2010); fewer microglia and parvalbumin interneurons (Lee et al., 2019); and a reduction in white matter PDGFRα^+^/NG2^+^ OPCs at early postnatal stages (Pacey et al., 2013). It is therefore plausible that Fmrp could regulate both the balanced production of neuronal and glial cells from a common progenitor domain and the subsequent differentiation of glial precursors into mature OLs.

We find that *fmr1* mutant zebrafish generate excess oligodendrocyte lineage cells (OLCs) throughout embryonic and larval development. Coupled with reduced motor neuron formation and Shh signaling, these data show a vital role for Fmrp in glial cell specification in embryonic development. In addition, we show that Fmrp promotes OL differentiation and limits the density of OPCs during subsequent larval stages. Taken together, Fmrp plays important roles in restricting glial precursor formation and driving myelin programming, two critical cellular programs in early neurodevelopment.

## Materials and Methods

### Zebrafish lines and husbandry

The Institutional Animal Care and Use Committee at the University of Colorado School of Medicine approved all animal work, which is in compliance with US National Research Council’s Guide for the Care and Use of Laboratory Animals, the US Public Health Service’s Policy on Humane Care and Use of Laboratory Animals, and Guide for the Care and Use of Laboratory Animals. Larvae were raised at 28.5°C in embryo medium and staged as hours or days post fertilization (hpf/dpf) according to morphological criteria (Kimmel et al., 1995). Zebrafish lines used in this study included *fmr1^hu2787^* (den Broeder et al., 2009), *Tg(myrf:mScarlet)^co66^* (this paper), and *Tg(olig2:EGFP)^vu12^* (Shin et al., 2003). Genotyping for *fmr1^hu2787^* was performed as previously described (Ng et al., 2013).

### Imaging and analysis

We acquired images on a Zeiss LSM 880 or a Zeiss CellObserver SD 25 spinning disk confocal system (Carl Zeiss). Images were captured with Zen software (Carl Zeiss), then processed and analyzed using Fiji/ImageJ or Zen Blue (Carl Zeiss).

### Immunohistochemistry

Larvae were fixed at indicated time points in 4% paraformaldehyde/1xPBS, rocking O/N at 4°C. Larvae were rinsed in 0.1%Triton/1xPBS (PBSTx), then embedded in 1.5% agar/30% sucrose and immersed in 30% sucrose O/N. Blocks were frozen on dry ice and 20 μm transverse sections were taken with a cryostat microtome and collected on polarized slides. Slides were mounted in Sequenza racks (Thermo Scientific), washed 3×5 minutes in PBSTx, blocked 1 hour in 2% goat serum/2% bovine serum albumin/PBSTx and then placed in primary antibody (in block) O/N: rabbit α-Sox10 (1:500; Park et al., 2005); mouse α-Islet (1:500; Developmental Studies Hybridoma Bank, AB2314683). Sections were washed 1.5 hours in PBSTx, and then incubated 2 hours at RT in secondary antibody (1:250; in block): AlexaFluor 488 goat α-rabbit (abcam, ab150077) and AlexaFluor 568 goat α-mouse (abcam, ab175473). Sections were washed for 1 hour in PBSTx, incubated with DAPI (1:2000 in PBSTx) for 5 minutes, washed 3×5 minutes in PBSTx, then mounted in Vectashield (Vector Laboratories, H-1000-10).

### *Fluorescent* in situ *RNA hybridization*

Fluorescent *in situ* RNA hybridization (FISH) was performed with the RNAScope Multiplex Fluorescent V2 Assay Kit (Advanced Cell Diagnostics). Embryos and larvae at indicated timepoints were fixed in 4% paraformaldehyde/1xPBS, rocking O/N at 4°C. Samples were then embedded in 1.5% agar/30% sucrose and immersed in 30% sucrose O/N. Blocks were frozen on dry ice and 12 μm transverse sections were taken with a cryostat microtome and collected on polarized slides. FISH was performed as per the manufacturer’s protocol with the following modifications: slides were covered with Parafilm for all 40°C incubations to maintain moisture and disperse reagents across sections. Probes for zebrafish *olig2-C1, nkx2.2-C2, ptch2-C3, sox10-C1, myrf-C2, cspg4-C3, neurog1-C2*, and *boc-C2* (1:50 dilution) were designed and synthesized by the manufacturer. Transcripts were labeled with the Opal 7 Kit (Perkin Elmer, NEL797001KT): Opal 520 (1:1500), Opal 570 (1:500), and Opal 650 (1:1500).

### Quantification and Statistical Analysis

For all cell counts, the investigator was blind to genotype. For IHC (Figs. 1, 3), DAPI and Sox10/Islet channels were used to confirm cell number in a given section. For FISH, (Fig. 4), DAPI *sox10, myrf, cspg4* were used to confirm cell number in acquired z-stacks. In experiments with transgenic larvae (Fig. 5), all quantification includes transgenic larvae expressing *myrf*:mScarlet, though not all larvae co-expressed *olig2*:EGFP.

**Figure 1.**
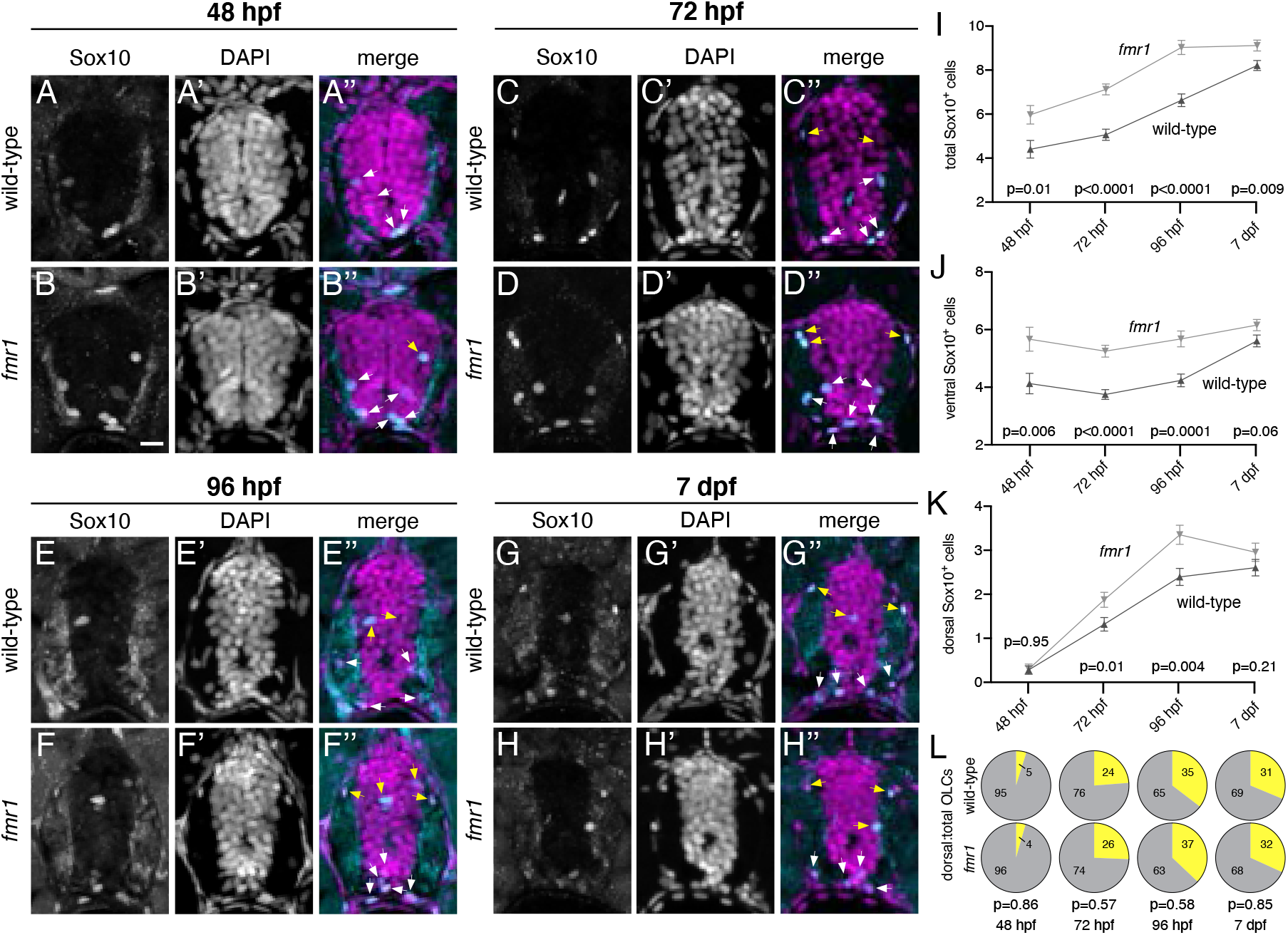
Fmrp restricts the production of OLCs. Representative images of trunk spinal cord transverse sections from wild-type and *fmr1* mutant embryos and larvae processed to detect Sox10 antibody expression at progressive developmental stages. Wild-type (A) and *fmr1* (B) embryos at 48 hpf (n^WT^=11 embryos, 32 sections; n^*fmr1*^=10 embryos, 30 sections). Wild-type (C) and *fmr1* (D) larvae at 72 hpf (n^WT^=10 larvae, 46 sections; n^*fmr1*^=10 larvae, 47 sections). Wild-type (E) and *fmr1* (F) larvae 96 hpf (n^WT^=12 larvae, 38 sections; n^*fmr1*^=12 larve, 37 sections). Wild-type (G) and *fmr1* (H) larvae at 7 dpf (n^WT^=11 larvae, 43 sections; n^*fmr1*^=13 larvae, 42 sections). White arrowheads indicate Sox10^+^ OLCs in the ventral half of the cord. Yellow arrowheads indicate OLCs in the dorsal half of the cord. Graphs depict total OLCs (I), ventrally located OLCs (J), dorsally located OLCs (K), and the ratio of dorsal to total OLCs (L) at each developmental stage (dorsal OLCs in yellow; total OLCs in gray). Significance determined by unpaired t tests (total OLCs; 7 dpf dorsal OPCs; 72hpf, 96hpf, 7dpf ratios) and Mann-Whitney tests (all others). Scale bars = 10 μm.

### Fluorescent RNA in situ hybridization quantification

FISH puncta were quantified from z-projections collected at identical exposures with an adapted ImageJ custom script created by Karlie Fedder, University of Colorado, Department of Pediatrics, https://github.com/rebeccaorourke-cu/Prdm8-regulates-pMN-progenitor-specification. First, ten z-intervals of 0.5 μm depth were maximum z-projected and background was subtracted with a 2-rolling ball. Next, the image was thresholded by taking two standard deviations above the mean fluorescence intensity. A region of interest was drawn around the spinal cord and puncta were analyzed using the “Analyze Particles” feature with a size of 0.01-Infinity and circularity of 0.00-1.00. All thresholded puncta were inspected to ensure single molecules were selected. Puncta with an area of only 1 pixel were removed from the dataset. Data for each embryo was collected from a minimum of three consecutives trunk spinal cord sections and n represents the average number of puncta in a region of interest per section.

### Statistics

All statistics were performed in Graphpad Prism (version 9). Normality was assessed with a D’Agostino and Pearson omnibus test. For two groups, unpaired comparisons were made using either unpaired two-tailed t tests (for normal distributions) or Mann-Whitney tests (abnormal distributions).

## Results

### Fmrp restricts the production of oligodendrocyte lineage cells

To examine the impact of Fmrp on gliogenesis in the developing spinal cord we used immunohistochemistry for Sox10 – a canonical marker of the oligodendrocyte lineage - on transverse sections of wild-type and *fmr1^hu2787^* mutant larvae at progressive embryonic and larval stages. Hereafter, we use oligodendrocyte lineage cell (OLC) to denote Sox10-expressing cells in the central nervous system. Our results show surplus OLCs in the absence of Fmrp, beginning with a ~26% increase in *fmr1* mutant embryos at 48 hours post-fertilization (hpf; Fig. 1A,B,I). At 72 hpf, there was a ~29% increase in OLCs in *fmr1* larvae (Fig. C,D,I), and a comparable ~27% increase at 96 hpf (Fig. 1E,F,I). Finally, at a late larval stage (7 days post-fertilization; dpf) there was a less pronounced ~10% increase in *fmr1* mutants compared to wild-type (Fig. 1 G,H,I). These results suggest that FMRP restricts the production of OLCs during early spinal cord development. Many OLCs generated from the pMN remain in the ventral cord, proximal to the prominent ventral axonal tracts. However, other OLCs migrate toward the myelinated tracts in the dorsal spinal cord (yellow arrowheads, Fig. 1). We next quantified dorsal and ventral OLCs to examine whether Fmrp regulates the regional distribution of cells. At 48 hpf, *fmr1* embryos had excess OLCs in the ventral cord compared to wild-type embryos (Fig. 1J), before the onset of most dorsal migration (see Fig. 2). By larval stages *fmr1* mutants had more OLCs in both the ventral and dorsal halves of the cord (Fig. 1J,K), such that the regional distribution OLCs was actually quite similar to wild-type controls throughout our developmental timeline (Fig. 1L). In short, these data indicate that Fmrp regulates the number of cells specified for OLC fate but not their proportional distribution between ventral and dorsal myelinated tracts of the spinal cord.

**Figure 2.**
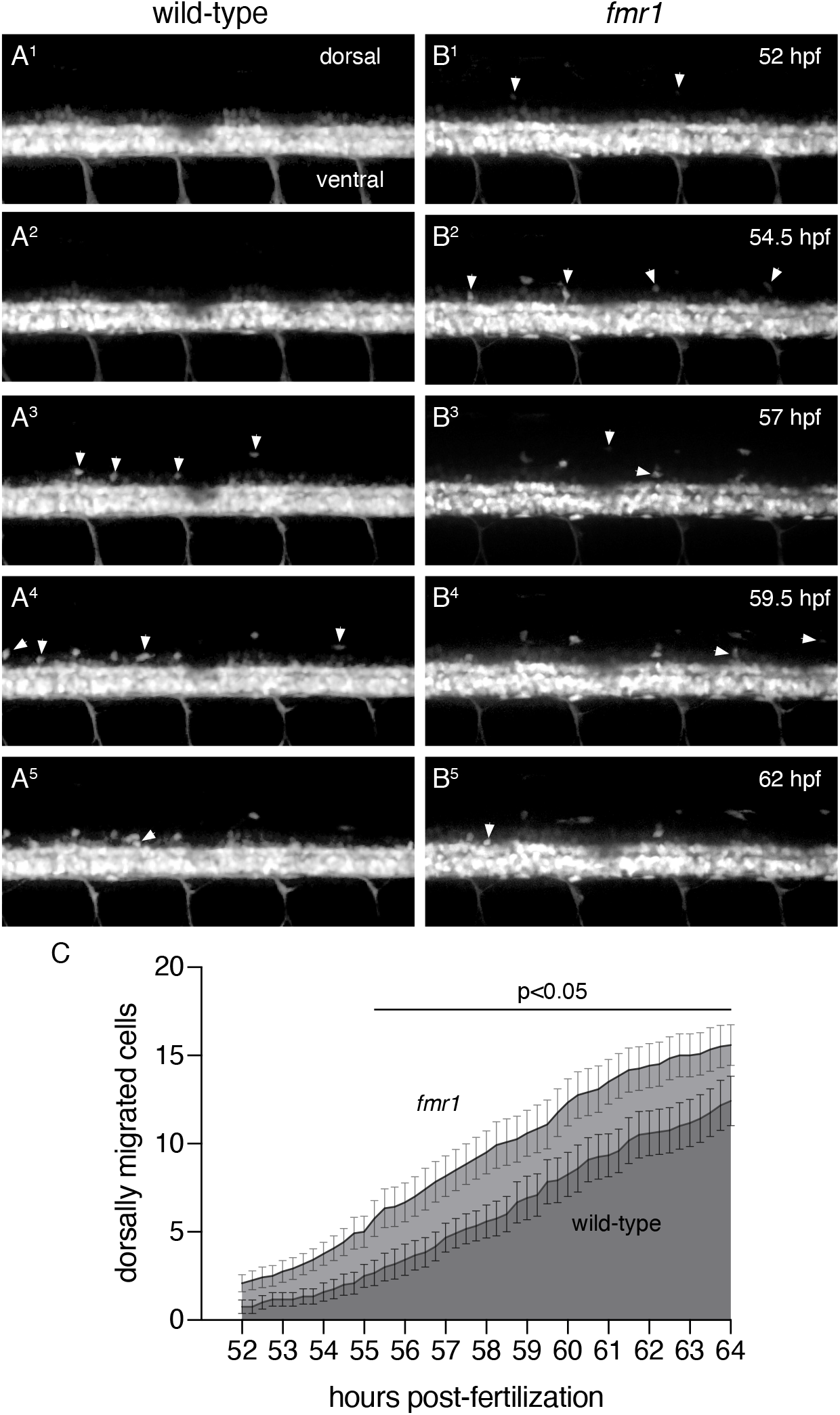
*fmr1* larvae show premature and excessive dorsal migration of OLCs. Representative lateral trunk images of live transgenic wild-type (A^1^-A^5^) and *fmr1* (B^1^-B^5^) embryos expressing *olig2:EGFP* at 2.5 hour intervals, beginning at 52 hpf. White arrowheads indicate OPCs that have migrated from the ventral spinal cord. (C) Graph depicts the average number of OPCs that have migrated from the ventral cord in *fmr1* (light gray) and wild-type (dark gray) at 15 minute increments. Significance determined by unpaired t tests at each time point. n=12 larvae for each genotype. Scale bar = 20 μm.

We next used time-lapse microscopy to better understand the timeline of dorsal OLC migration in both wild-type and *fmr1* mutants. We captured images of living transgenic larvae expressing *olig2:EGFP* – a marker of cells derived from the pMN domain, including OLCs (Shin et al., 2003) - at the onset of dorsal OLC migration, recording lateral threedimensional z-stacks through the cord every 15 minutes from 52 to 64 hpf (Fig. 2A,B). We then analyzed the timing of OLC emergence from the dense *olig2:*EGFP^+^ ventral domain. This developmental window was based on our Sox10 immunohistochemistry timeline, which showed that very few OLCs had migrated dorsally at 48 hpf (Fig.1). It is also important to note wild-type and *fmr1* larvae, indicating that we captured the bulk of migration in most larvae. During much of the subsequent timeline, that migrating *olig2^+^* cells are OLCs, as pMN-derived motor neurons do not leave the ventral cord (see Fig. 3). At the onset of imaging at 52 hpf, there were very few cells in the dorsal spinal cord in both specifically 55 to 64 hpf, there was premature and excessive dorsal migration of OLCs in *fmr1* larvae compared to wild-type controls (Fig. 2C). Taken together with our previous data showing excess OLCs in *fmr1* (Fig. 1), these results raise the possibility that Fmrp restricts OLC formation and suggests that the precocious dorsal migration of cells in *fmr1* larvae is likely due to increased cell density.

**Figure 3.**
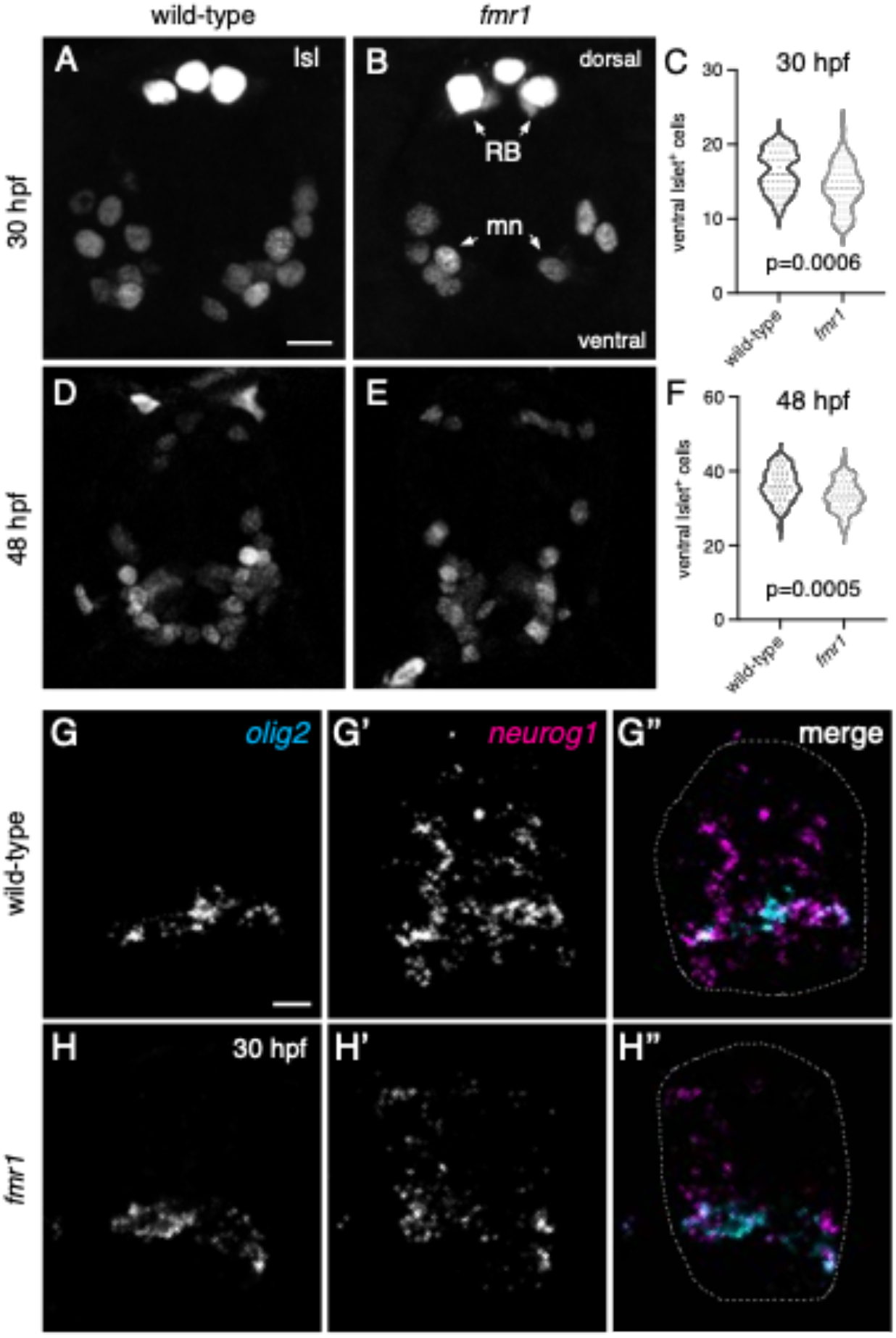
Fmrp promotes motor neuron formation. Representative images of trunk spinal cord transverse sections processed to detect expression of Islet protein (IsI) In wild-type (A) and *fmr1* (B) embryos at 30 hpf. (C) Graph comparing the quantity of IsI^+^ motor neurons In wild-type and *fmr1* embryos at 30 hpf (n^WT^=15 embryos. 55 sections; n^*fmr1*^=15 embryos, 57 sections). Representative Images of Is^+^ motor neurons In wild-type (D) and *fmr1* (E) embryos at 48 hpf. (C) Graph comparing the quantity of IsI^+^ motor neurons In wild-type and *fmr1* larvae at 48 hpf (n^wT^=15 embryos, 55 sections: n^*fmr1*^=15 embryos. 58 sections). RB=Rohon Beard neurons; mn=motoneurons. Significance determined by Mann-Whitney test (30 hpf) and unpaired t test (48 hpf). Representative Images of transverse spinal cord sections processed to detect expression of *olig2 and neurogenin1* (*neurog1*) mRNA In wild-type (G) and *fmr1* mutant embryos (H) at 30 hpf (n= 5 embryos, 25 sections each genotype). Spinal cord in G”,H” outlined in dashed line. Scale bars = 10 μm.

### Fmrp promotes motor neuron formation

The excess OLCs noted in *fmr1* mutants led us to examine whether motor neuron formation was also affected by the loss of Fmrp, as pMN progenitor cells give rise to both motor neurons and OPCs during embryonic development (Park et al., 2002). We therefore quantified Islet^+^ (Isl) cells - an early marker of motor neurons (Ericson et al., 1992) - in transverse sections of wild-type and *fmr1* mutant embryos (Fig. 3). At 30 hpf, there was a ~13% reduction in Isl^+^ motor neurons in *fmr1* embryos compared to wild-type control (Fig. 3A-C). At 48 hpf, there was a less pronounced ~8% reduction in *fmr1* embryos (Fig. 3D-F). These results suggest that Fmrp also promotes neurogenesis in the pMN progenitor domain.

The excess OLCs and concomitant reduction in motor neurons in *fmr1* mutants could indicate that Fmrp regulates cell formation in the pMN by promoting a pro-neurogenic gene expression program. To test this, we used fluorescent RNA in situ hybridization to examine proneuronal gene expression in wild-type and *fmr1* embryos at 30 hpf. *neurogenin1 (neurog1*; Blader et al., 1997) expression was prominent throughout the spinal cord of wild-type embryos, including considerable overlap with *olig2* in the pMN domain (Fig. 3G). In contrast, there was a sharp reduction in *neurog1* expression in *fmr1* mutants as compared to control embryos (Fig. 3H). There were no obvious changes in *olig2* expression in *fmr1* embryos, which suggests that Fmrp is not required for progenitor cell formation. Taken together, these data indicate that FMRP promotes proneuronal gene expression and motor neurogenesis in the developing embryonic spinal cord, at a developmental stage that coincides with increased OLC formation in *fmr1* mutants (Fig. 1).

### Fmrp promotes oligodendrocyte differentiation

The reduced white matter in infants with Fragile X syndrome (Swanson et al., 2018) and deficient myelination in FXS model organisms (Pacey et al., 2013; Doll et al., 2019) could reflect a requirement for Fmrp in the differentiation of OLCs into myelinating oligodendrocytes. We examined this question in late embryonic and larval stages using three different methods. First, we used fluorescent RNA in situ hybridization to examine expression of three genes associated with progressive developmental stages of the OL lineage: *olig2*, a marker of pMN cells and OLCs; *sox10* an OLC marker; and *myelin regulatory factor* (*myrf*), a gene expressed in differentiating OLs (Hornig et al., 2013). At 48 hpf, there was ~21% increase in total *sox10^+^* OLCs in *fmr1* mutants compared to wild-type (Fig 4A,B,C), a result comparable to our immunohistochemical results using Sox10 antibody (Fig. 1). Although the quantity of *sox10^+^* cells co-expressing *myrf* was not significantly different between genotypes (Fig 4D, pink outlines in 4A’’’,4B’’’), there was a ~33% increase in *sox10^+^* cells that did not express *myrf* in *fmr1* mutants (Fig. 4E, yellow outlines in 4A’”,4B’”). When taken as a ratio of total *sox10*^+^ OLCs, 66% of wildtype cells expressed *myrf*, but only 59% of cells in *fmr1* mutants coexpressed both genes (Fig. 4F). These data suggest that proportionately fewer OLCs are undergoing differentiation in *fmr1* mutants at this late embryonic stage, despite the preponderance of total OLCs.

**Figure 4.**
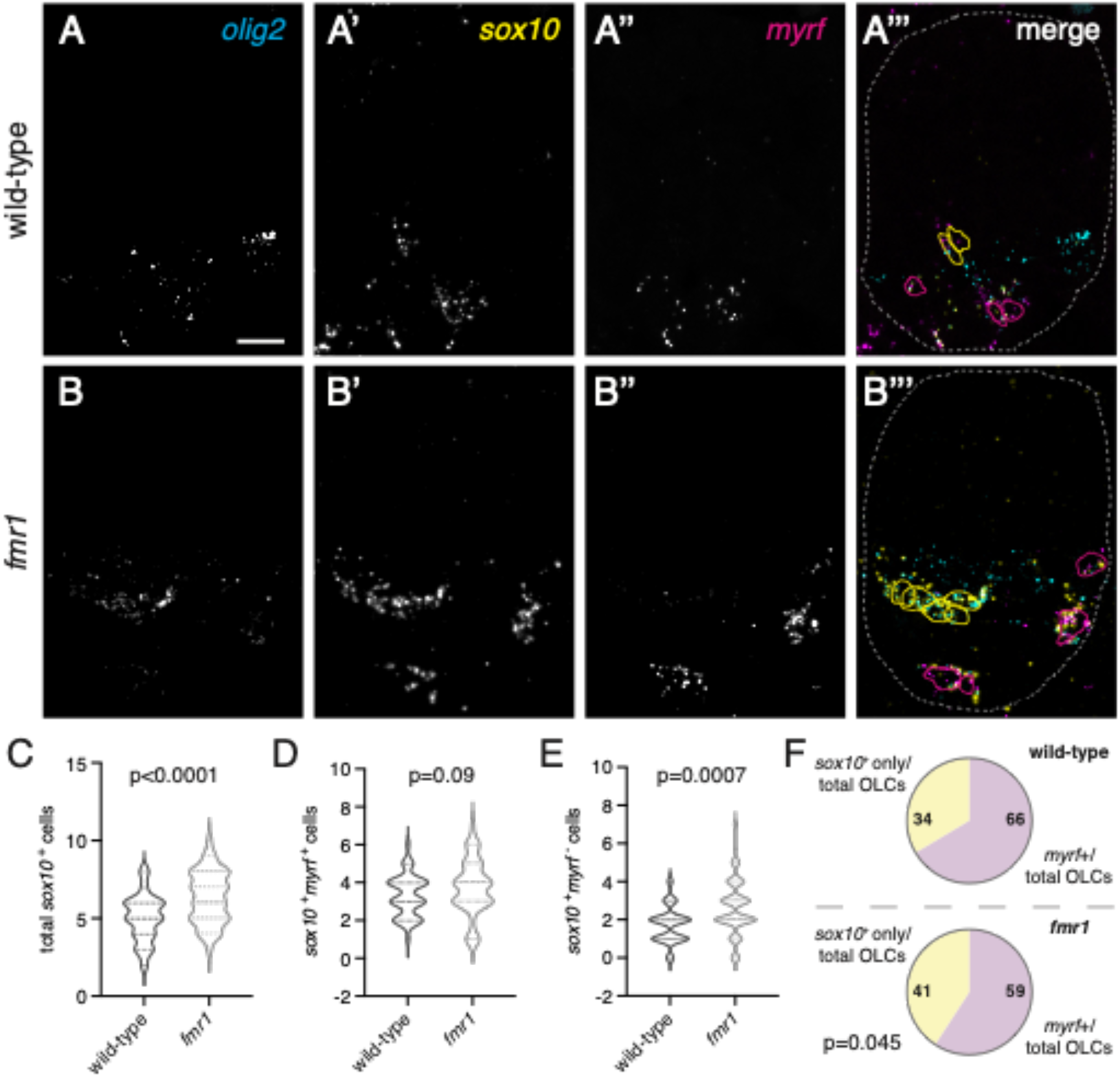
Fmrp promotes oligodendrocyte differentiation In the embryonic spinal cord. Representative images of trunk spinal cord sections from wild-type (A-A’”) and *fmr1* mutant embryos (B-B”’) at 30 hpf, processed to detect expression of specific mRNAs using fluorescent in situ hybridization, including pMN descendants (*olig2*; A,B), oligodendrocyte lineage cells (*sox10*; A’.B’), and differentiating oligodendrocytes (*myrf*: A”,B”). Cell outlines in A”’ and B”’ mark *olig2^+^sox10^+^myrf* cells (yellow) and *ollg2^+^so×10^+^myrf^+^* cells (magenta), dashed lines represent the edges of the spinal cord. Graphs comparing the average total OLCs (C), *myrf^+^* OLCs (D), and *myrf* OLCs (E) in wild-type (n=9, 45 sections) and *fmr1* (n=10, 48 sections). (F) The relative ratio of *myrf^+^* cells to total OLCs in wild-type and *fmr1* mutants. Significance determined with unpaired t tests. Scale bar = 10 μm.

As an independent method to assess the timing of OL differentiation in live animals, we next quantified differentiating OLs in subsequent larval stages using stable transgenic lines. We crossed *Tg(olig2:EGFP)^vu12^*, a reporter of pMN lineage cells (Fig. 5A,B,C,D), with *Tg(myrf:mScarlet)^co66^*, a reporter of differentiating oligodendrocytes (Fig. 5,A’,B’,C’,D’). At 72 hpf, there was a ~44% reduction in total *myrf*^+^ cells in *fmr1* larvae compared to wild-type (Fig. 5E), though it is important to note that this approach may be less sensitive than FISH (Fig. 4) due to the folding time of the mScarlet fluorophore. At 96 hpf, *fmr1* larvae had a less pronounced ~11% reduction in *myrf^+^* cells compared to controls (Fig. 5E), further indicating a delay in OL differentiation. Although *fmr1* larvae had surplus OLCs in the dorsal spinal cord compared to wild-type controls (Fig. 5F) - consistent with our time-lapse imaging (Fig. 2) - far fewer of these migrated cells expressed *myrf* (Fig. 5G). Indeed, at these larval stages *fmr1* mutants had a reduced ratio of differentiating *myrf*^+^ cells to total OLCs in the dorsal spinal cord compared to wild-type (Fig. 5H). These results further validate the reduced OL differentiation seen in *fmr1* mutants and suggest OL differentiation is delayed in the absence of Fmrp.

**Figure 5.**
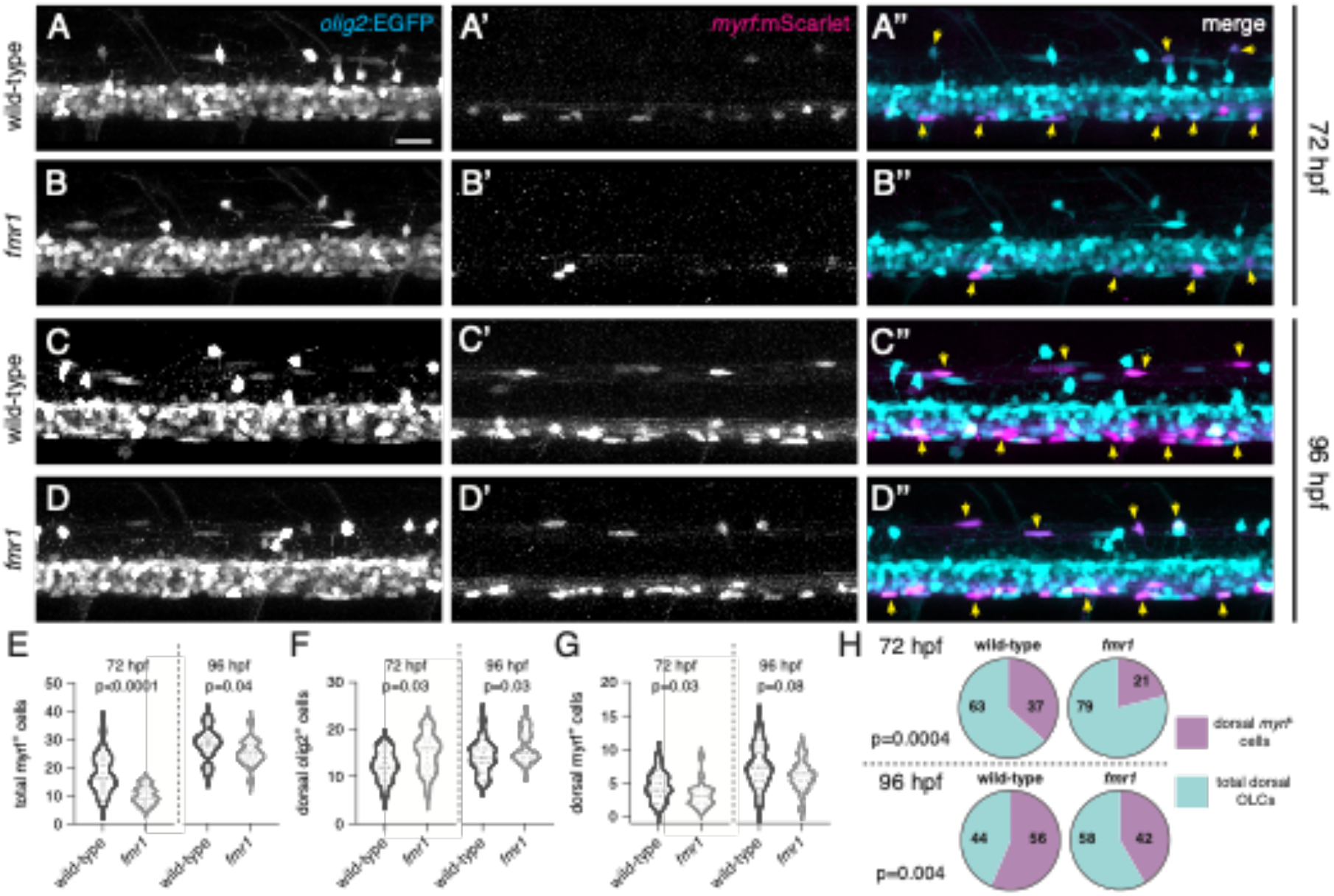
Fmrp regulates the timing of OL differentiation. Representative lateral Images of live transgenic wild-type and *fmr1* mutant larvae at 72 hpf (A-B) and 96 hpf (C-D) stably expressing *olig2:EGFP* (pMN-derlved cells) and myrf.mScarlet (differentiating oligodendrocytes). Yellow arrowheads Indicate *olig2^+^myrf*^+^ cells. Graphs comparing the quantity of total *myrf*^+^ cells (E; n^*72hpf*^=20 larvae each, n^*96hpf*^=24^WT^,23^*fmr1*^), dorsal OLCs (F; n^*72hpf*^=19 each, n^*96hpf*^=20^WT^,23^*fmr1*^), dorsal *myrf*^+^ cells (G; n^*72hpf*^=20 each. n^*96hpf*^=24^WT^,23^fmr1^), and the ratio of differentiating dorsal OLCs (H; n^*72hpf*^=19 each, n^*96hpf*^=20^WT^,22^*fmr1*^). Significance determined by a Mann-Whitney tests (G, 72 hpf) and unpaired t tests (all others). Scale bar = 20 μm.

The above data suggest that a smaller proportion of OPCs undergo differentiation into myelinating OLs in *fmr1* mutant larvae than in wild-type and thus OPCs may retain a longer-term precursor identity in the absence of Fmrp. To test this possibility directly, we again used fluorescent RNA in situ hybridization at a late larval stage to detect expression of *chondroitin sulfate proteoglycan 4 (cspg4)*, which marks OPCs in larval and adult stages, alongside *sox10* and *myrf* (Fig. 6). At 6 dpf, *fmr1* larvae had ~40% more *cspg4^+^* cells than wild-type controls (Fig. 6A,B,C) and ~42% of total OLCs in *fmr1* larvae expressed *cspg4,* compared to just ~34% of OLCs in wild-type (Fig. 6F), indicating that more OLCs retain a precursor identity in the absence of Fmrp. In addition, there was no change in the quantity of *myrf*^+^ differentiating OLCs in *fmr1* larvae compared to wild-type at this late larval stage (Fig. 6A’,B’,D), which further suggests that Fmrp regulates the timing of differentiation, as there were fewer differentiating OLs in *fmr1* mutants at earlier larval stages (Fig. 5). Finally, *fmr1* larvae had ~23% more total *sox10*^+^ OLCs than wildtype at 6 dpf (Fig. 6A”,B”,E), which was in line with Sox10 immunohistochemistry at late larval stages (Fig. 1). Taken together, these data suggest that Fmrp regulates both the timing of OL differentiation and the conversion of OPCs to OLs, such that *fmr1* mutants have more OPCs at late larval stages.

**Figure 6.**
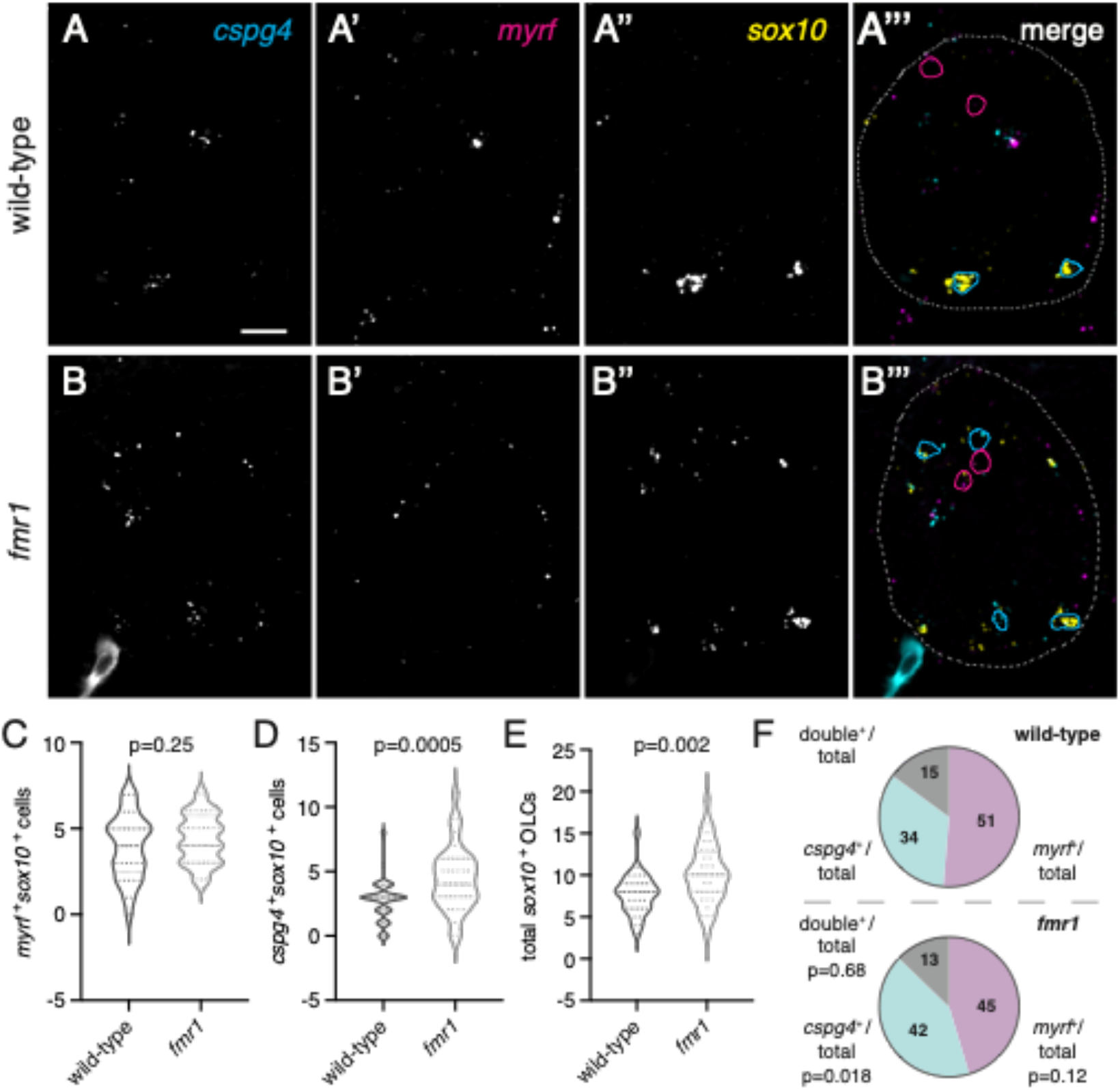
*fmr1* mutant larvae have excess OPCs. Representative Images of transverse trunk spinal cord sections of wild-type (A-A’”) and *fmr1* mutant (B-B’“) larvae processed to detect expression of *cspg4a, myrf*, and *sox10* at 6 dpf via fluorescent in situ RNA hybridization. (A’”,B”‘) *sox10^+^myrf^+^cspg4^+^* cells are outlined in magenta; *cspg4^+^* cells not expressing *myrf* in cyan; spinal cord boundaries in dashed lines. Graphs comparing *myrf^+^* OLCs (C). *cspg4^+^* OLCs (D). and total *sox10^+^* cells In wild-type (n=8 larvae. 33 sections) and *fmr1* mutants (n=8 larvae, 36 sections). (F) Relative ratios of OLCs expressing *myrf, cspg4*, or co-expressing both transcripts. Significance determined by unpaired t tests (total *sox10^+^, myrf^+^* comparisons) and Mann-Whitney tests (double^+^ ratio, *cspg4^+^* compansons). Scale bar = 10 μm.

### fmr1 *mutants have abnormally low levels of Shh signaling in the developing spinal cord*

The Sonic hedgehog (Shh) signaling pathway is essential for the dorsoventral patterning of the spinal cord (Briscoe et al., 2000; Dessaud et al., 2007) and a transient increase in Shh signaling is necessary for OPC formation (Al Oustah et al., 2014; Scott et al., 2020). Moreover, OL formation appears to require slightly higher levels of Shh signaling than OPCs, as pharmacological inhibition of Shh leads to more OPCs and fewer OLs and overexpression of Shh shifts the balance toward OL fate (Ravanelli et al., 2018). To test whether altered populations of OLCs and motor neurons in *fmr1* mutants are associated with changes in Shh signaling, we used fluorescent RNA in situ hybridization at early embryonic stages to examine the expression of genes marking ventral progenitor domains and Shh signaling: *olig2*, a marker of the pMN progenitor domain; *nkx2.2*, a marker of the p3 progenitor domain; *ptch2*, a positive transcriptional target of Shh (Concordet et al., 1996); and *boc*, a Shh coreceptor and negative transcriptional target of Shh (Tenzen et al., 2006; Kearns et al., 2021). *fmr1* mutant embryos had no obvious changes relative to wild-type in the expression of *olig2* at 24 or 30 hpf (Fig. 7A,B,E,F) or *nkx2.2* at 30 hpf (Fig. 7E’,F’), which again suggests a general maintenance of spinal cord progenitor boundaries in *fmr1* mutants. However, *fmr1* embryos had a clear reduction in *ptch2* expression at both 24 hpf (Fig. 7A”,B”) and 30 hpf (Fig. 7E”,F”), and *boc* expression was increased at 24 hpf compared with wild-type embryos (Fig. 7A’,B’). We quantified the total number of *boc^+^* puncta at 24 hpf and *ptch2*^+^ puncta at 30 hpf in individual transverse trunk spinal cord sections (Fig. 7C,G) and normalized this data to the relative size of the spinal cord. At 24 hpf, *fmr1* mutants had ~19% more *boc^+^* puncta than wild-type (Fig. 7D). In addition, *fmr1* mutants had a ~26% decrease in *ptch2^+^* puncta compared to wild-type at 30 hpf (Fig. 7H). Taken together, these results raise the possibility that Fmrp regulates OLC formation, at least in part, by shaping or maintaining the Shh signaling gradient.

**Figure 7.**
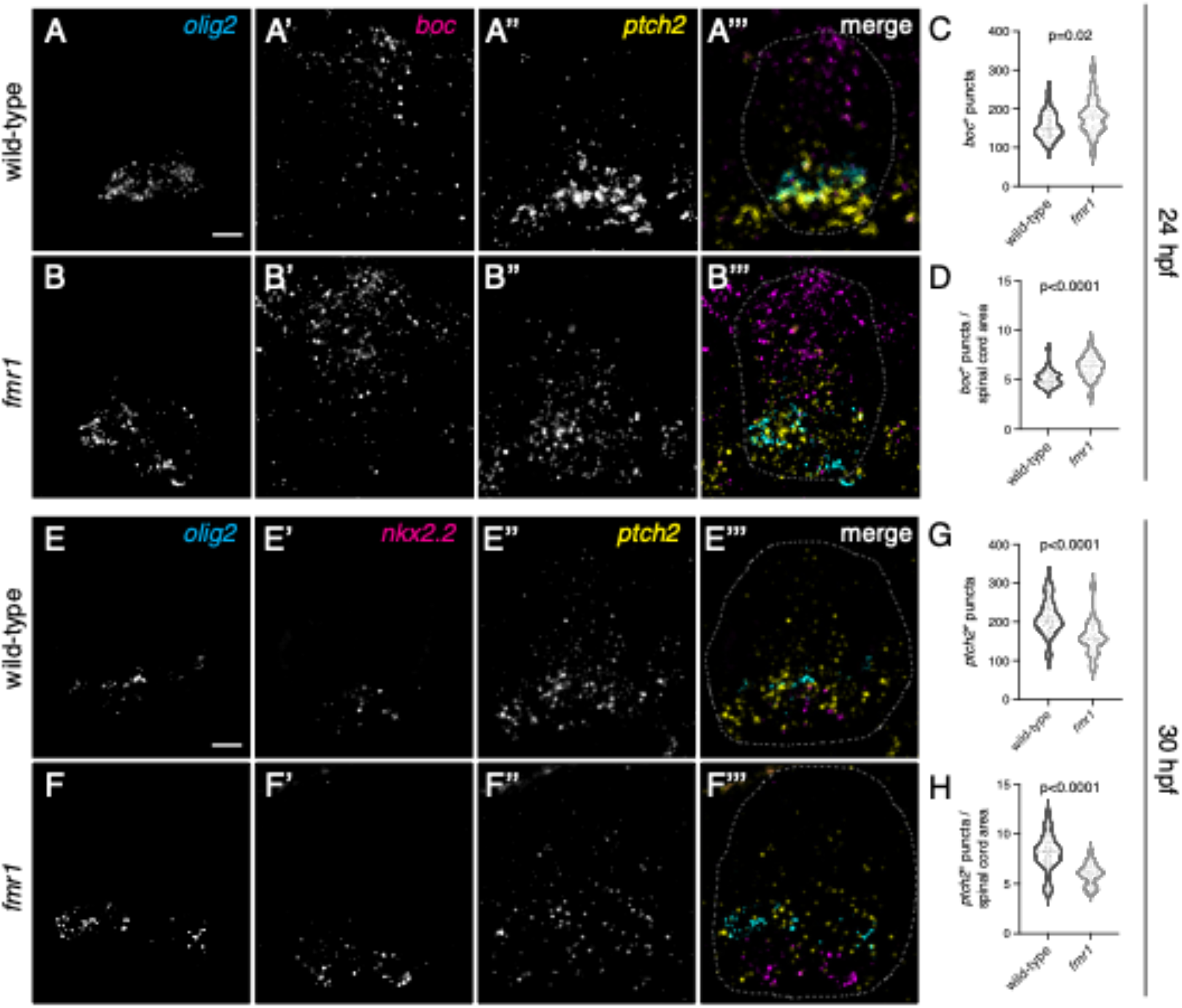
*fmr1* mutants have abnormally low levels of Shh signaling In the developing spinal cord. Representative images of transverse trunk spinal cord sections from 24 hpf wild-type (A-A’”) and *fmr1* mutants (B-B’”) processed to detect expression of *olig2, boc*. and *ptch2* using fluorescent in situ RNA hybridization. Graphs comparing the average number of *boc^+^* puncta per section (C; unpaired t test), and average number of *boc^+^* puncta per unit area of spinal cord (D; Mann-Whitney test) in wild-type (n=6 embryos, 30 sections) and *fmr1* mutants (n=6 embryos, 30 sections). Representative images of transverse trunk spinal cord sections from 30 hpf wild-type (E-E’”) and *fmr1* mutants (F-F’”) processed to detect expression of *olig2, nkx2.2*, and *ptch* using fluorescent in situ RNA hybndization. Graphs comparing the average number of *ptch^+^* puncta per section (G; Mann-Whitney test), and average number of *ptch^+^* puncta per unit area (H; unpaired t test) in wild-type (n=7 embryos, 35 sections) and *fmr1* mutants (n=6 embryos, 26 sections). Scale bars = 10 μm.

## Discussion

The proportionate balance and integration of a wide variety of cell types is crucial for neural circuit formation and function. For example, altered quantities of excitatory and inhibitory neurons in the absence of FMRP (Tervonen et al., 2009; Lee et al., 2019) could directly contribute to the hyperexcitation theory of FXS (Gibson et al., 2008; Paluszkiewicz et al., 2011; Contractor et al., 2015). Imbalanced excitation (E) and inhibition (I) may contribute to debilitating symptoms in ASDs at large, such as hypersensitivity to sensory stimuli (Sapey-Triomphe et al., 2019). Importantly, infants with FXS have diminished white matter profiles (Swanson et al., 2018), and OLs autonomously require Fmrp for myelin sheath growth (Doll et al., 2019), which implicate glia in the disease state. In another interesting parallel, social isolation during an early critical period, leads to dramatic decreases in myelination (Makinodan et al., 2012). Oligodendrocyte lineage cells dramatically influence neural circuits, as myelin plasticity is required for many learning and memory paradigms (Mckenzie et al., 2014; Pan et al., 2020; Wang et al., 2020), and OPCs can also directly modulate synaptic transmission (Sakry et al., 2014). Finally, the fact that OLs ensheathe both inhibitory and excitatory axons with myelin (Micheva et al., 2016) further indicates that oligodendroglia can directly influence the balance of excitation and inhibition in the nervous system.

Our current study shows that Fmrp promotes OL differentiation, which could help explain the poor myelin sheath growth in the OLs that do manage to differentiate in *fmr1* mutant zebrafish. This is perhaps not surprising given the catalogue of mRNA targets of FMRP associated with differentiation (Liu et al., 2018). The reduced OL differentiation seen in *fmr1* mutants could be linked to diminished Shh in early embryonic development (Fig. 7), as our lab previously showed that pharmacological inhibition of Shh impedes OLC differentiation, resulting in excess OPCs and fewer mature OLs (Ravanelli et al., 2018). Interestingly, *Fmr1-knockout* NPCs are also differentiation-impaired, leading to surplus progenitors and fewer mature neurons (Edens et al., 2019). In this context, FMRP acts as an m^6^A reader driving nuclear export of mRNA; m^6^A methylation also promotes OL differentiation and myelination (Xu et al., 2020). These studies therefore represent striking parallels to this study and our previous work (Doll et al., 2019). In addition, overexpression of the NG2 proteoglycan intracellular domain in OPCs leads to reduced FMRP and increased translation (Nayak et al., 2018), which suggests that Fmrp may repress translation of mRNAs maintaining OPC fate. Taken together, our current results (Figs. 4-6) provide ample evidence that Fmrp promotes differentiation in the OL lineage.

Many questions remain in regard to how Fmrp regulates gliogenesis and the fate decision between OPCs and OLs. Does excess OPC production in *fmr1* mutants reflect a feedback mechanism linking diminished OL differentiation with prolonged gliogenesis? Are OPCs in *fmr1* larvae playing specialized roles in developing neural circuits lacking Fmrp? Persistent *cspg4*-expressing OPCs (NG2 glia) are noted throughout adulthood in wild-type animals (Dawson et al., 2003b; Simon et al., 2011) and are not simply a reserve pool to replace lost oligodendrocytes: NG2 glia are directly responsive to neuronal activity (Bergles et al., 2000) and can actively modulate synapses (Sakry et al., 2014). Interestingly, although there are actually fewer deep white matter OPCs in a mouse model of FXS at early postnatal stages (Pacey et al., 2013), both mouse and zebrafish models show deficient myelination (Pacey et al., 2013; Doll et al., 2019). This may indicate region-specific roles for the Fmrp in gliogenesis but a common requirement in OL differentiation. Nevertheless, the increase in persistent spinal *cspg4^+^* OPCs in *fmr1* mutants at late larval stages leave intriguing possibilities, especially when taking into account the overall increase in total OLCs (Fig. 6). Interestingly, OPCs in the zebrafish spinal cord are heterogenous, including one subgroup more prone to differentiate and another subset that displays increased calcium and process dynamics (Marisca et al., 2020). In the future we plan to characterize the impact of Fmrp loss on the morphology and excitability of both spinal neurons and OPCs.

Our work also links increased OPC density in *fmr1* mutants with precocious dorsal migration in the developing spinal cord (Figs. 1,2). OPC migration is heavily influenced by neighboring OPCs, which maintain spacing through contact-mediated repulsion (Kirby et al., 2006; Hughes et al., 2013). We speculate that the excess and premature dorsal migration of OPCs in *fmr1* mutants may stem from a similar repulsive mechanism, as we also noted clusters of Sox10^+^ OPCs in the ventral spinal cord during late embryogenesis (48 hpf), just prior to the onset of dorsal migration (Fig. 1B). Although we cannot completely rule out a role for Fmrp in migration, the regional distribution of OLCs in the spinal cord was remarkably consistent between wild-type and *fmr1* mutants (Fig. 1L), which suggests that OPC migration is largely dependent on cell density.

FMRP is capable of regulating nearly every stage of the mRNA lifecycle, including nuclear shuttling (Edens et al., 2019), localization (Dictenberg et al., 2008; Pilaz et al., 2016), translation (Todd et al., 2003), and stability (Zalfa et al., 2007). Although we cannot yet determine the precise mRNA targets regulating gliogenesis and differentiation, our future work will identify the FMRP-bound mRNAs in defined stages of the OL lineage. We predict that FMRP requirements are stage specific, such that the catalog of mRNAs regulated in progenitor cells is largely distinct from the highly motile precursors, and also different from the mRNAs localized in distal OL processes driving myelin sheath growth (Doll et al., 2019). Here we show stage specific defects in the oligodendrocyte lineage of *fmr1* mutants: FMRP appears to restrict the formation of persistent OPCs, while promoting differentiation of mature OLs. Taken together, our work unveils a crucial role for FMRP in the OL lineage with the potential for widespread consequences on neural circuit formation and function.

## Author Contributions

C.D. and B.A. conceived the project. K.S. performed fluorescent in situ experiments and quantification. C.D. performed all additional experiments. C.D. analyzed all data. C.D. wrote and B.A. edited the manuscript.

## Acknowledgments

The authors would like to thank the members of the lab for insightful discussion. This work was supported by US National Institute of Health (NIH) grant R21 NS117886 to C.D, R21 NS110213 to C.D. and B.A., and R01 NS095679 and a gift from the Gates Frontiers Fund to B.A. The University of Colorado Anschutz Medical Campus Zebrafish Core Facility was supported by NIH grant P30 NS048154. The anti-Isl antibody, developed by T.M. Jessell and S. Brenner-Morton, was obtained from the Developmental Studies Hybridoma Bank, created by the NICHD of the NIH and maintained by The University of Iowa, Department of Biology, Iowa City, IA 52242.

## Data Availability

The data that support the findings of this study are available from the corresponding author upon reasonable request.

## References

Ascano M, Mukherjee N, Bandaru P, Miller JB, Nusbaum JD, Corcoran DL, Langlois C, Munschauer M, Dewell S, Hafner M, Williams Z, Ohler U, Tuschl T. 2012. FMRP targets distinct mRNA sequence elements to regulate protein expression. Nature 492:382–386.

Barone R, Fichera M, De Grandi M, Battaglia M, Lo Faro V, Mattina T, Rizzo R. 2017. Familial 18q12.2 deletion supports the role of RNA-binding protein CELF4 in autism spectrum disorders. Am J Med Genet Part A 173:1649–1655.

Bergles DE, Roberts JDB, Somogyi P, Jahr CE. 2000. Glutamatergic synapses on oligodendrocyte precursor cells in the hippocampus. Nature 405:187–191.

Blader P, Fischer N, Gradwohl G, Guillemot F, Strähle U. 1997. The activity of Neurogenin1 is controlled by local cues in the zebrafish embryo. Development 124:4557–4569.

Briscoe J, Pierani A, Jessell TM, Ericson J. 2000. A homeodomain protein code specifies progenitor cell identity and neuronal fate in the ventral neural tube. Cell 101:435–445.

den Broeder MJ, van der Linde H, Brouwer JR, Oostra BA, Willemsen R, Ketting RF. 2009. Generation and characterization of Fmr1 knockout zebrafish. PLoS One 4:2–7.

Concordet JP, Lewis KE, Moore JW, Goodrich L V., Johnson RL, Scott MP, Ingham PW. 1996. Spatial regulation of a zebrafish patched homologue reflects the roles of sonic hedgehog and protein kinase A in neural tube and somite patterning. Development 122:2835–2846.

Contractor A, Klyachko VA, Portera-Cailliau C. 2015. Altered Neuronal and Circuit Excitability in Fragile X Syndrome. Neuron 87:699–715.

Darnell JC, Van Driesche SJ, Zhang C, Hung KYS, Mele A, Fraser CE, Stone EF, Chen C, Fak JJ, Chi SW, Licatalosi DD, Richter JD, Darnell RB. 2011. FMRP stalls ribosomal translocation on mRNAs linked to synaptic function and autism. Cell 146:247–261.

Dawson MRL, Polito A, Levine JM, Reynolds R. 2003a. NG2-expressing glial progenitor cells: An abundant and widespread population of cycling cells in the adult rat CNS. Mol Cell Neurosci.

Dawson MRL, Polito A, Levine JM, Reynolds R. 2003b. NG2-expressing glial progenitor cells: an abundant and widespread population of cycling cells in the adult rat CNS. 24:476–488.

Dessaud E, Yang LL, Hill K, Cox B, Ulloa F, Ribeiro A, Mynett A, Novitch BG, Briscoe J. 2007. Interpretation of the sonic hedgehog morphogen gradient by a temporal adaptation mechanism. Nature 450:717–720.

Dictenberg JB, Swanger SA, Antar LN, Singer RH, Bassell GJ. 2008. A Direct Role for FMRP in Activity-Dependent Dendritic mRNA Transport Links Filopodial-Spine Morphogenesis to Fragile X Syndrome. Dev Cell 14:926–939.

Doll CA, Broadie K. 2015. Activity-dependent FMRP requirements in development of the neural circuitry of learning and memory. Development 142:1346–1356.

Doll CA, Vita DJ, Broadie K. 2017. Fragile X Mental Retardation Protein Requirements in Activity-Dependent Critical Period Neural Circuit Refinement. Curr Biol 27:2318–2330.e3.

Doll CA, Yergert KM, Appel BH. 2019. The RNA binding protein fragile X mental retardation protein promotes myelin sheath growth. Glia 68:495–508.

Edens BM, Vissers C, Su J, Arumugam S, Xu Z, Shi H, Miller N, Rojas Ringeling F, Ming G, He C, Song H, Ma YC. 2019. FMRP Modulates Neural Differentiation through m6A-Dependent mRNA Nuclear Export. Cell Rep 28:845–854.e5.

Ericson J, Thor S, Edlund T, Jessell T, Yamada T. 1992. Early stages of motor neuron differentiation revealed by expression of homeobox gene Islet-1. Science (80-) 256:1555–1560.

Gibson JR, Bartley AF, Hays SA, Huber KM. 2008. Imbalance of neocortical excitation and inhibition and altered UP states reflect network hyperexcitability in the mouse model of fragile X syndrome. J Neurophysiol 100:2615–2626.

Hornig J, Fröb F, Vogl MR, Hermans-Borgmeyer I, Tamm ER, Wegner M. 2013. The Transcription Factors Sox10 and Myrf Define an Essential Regulatory Network Module in Differentiating Oligodendrocytes. PLoS Genet 9.

Hughes EG, Kang SH, Fukaya M, Bergles DE. 2013. Oligodendrocyte progenitors balance growth with self-repulsion to achieve homeostasis in the adult brain. Nat Neurosci 16:668–676.

Kearns CA, Walker M, Ravanelli AM, Scott K, Arzbecker MR, Appel B. 2021. Zebrafish spinal cord oligodendrocyte formation requires boc function. bioRxiv:1–9.

Kimmel CB, Ballard WW, Kimmel SR, Ullmann B, Schilling TF. 1995. Stages of embryonic development of the zebrafish. Dev Dyn 203:253–310.

Kirby BB, Takada N, Latimer AJ, Shin J, Carney TJ, Kelsh RN, Appel B. 2006. In vivo time-lapse imaging shows dynamic oligodendrocyte progenitor behavior during zebrafish development. Nat Neurosci 9:1506–1511.

Lee FHF, Lai TKY, Su P, Liu F. 2019. Altered cortical Cytoarchitecture in the Fmr1 knockout mouse. Mol Brain 12:1–12.

Lee JA, Damianov A, Lin CH, Fontes M, Parikshak NN, Anderson ES, Geschwind DH, Black DL, Martin KC. 2016. Cytoplasmic Rbfox1 Regulates the Expression of Synaptic and Autism-Related Genes. Neuron 89:113–128.

Liu B, Li Y, Stackpole EE, Novak A, Gao Y, Zhao Y, Zhao X, Richter JD. 2018. Regulatory discrimination of mRNAs by FMRP controls mouse adult neural stem cell differentiation. Proc Natl Acad Sci U S A 115:E11397–E11405.

Luo Y, Shan G, Guo W, Smrt RD, Johnson EB, Li X, Pfeiffer RL, Szulwach KE, Duan R, Barkho BZ, Li W, Liu C, Jin P, Zhao X. 2010. Fragile X Mental Retardation Protein Regulates Proliferation and Differentiation of Adult Neural Stem/Progenitor Cells. PLoS Genet 6:e1000898.

Makinodan M, Rosen KM, Ito S, Corfas G. 2012. A critical period for social experience-dependent oligodendrocyte maturation and myelination. Science (80-) 337:1357–1360.

Marisca R, Hoche T, Agirre E, Hoodless LJ, Barkey W, Auer F, Castelo-Branco G, Czopka T.2020. Functionally distinct subgroups of oligodendrocyte precursor cells integrate neural activity and execute myelin formation. Nat Neurosci 23:363–374.

Mckenzie IA, Ohayon D, Li H, Faria JP De, Emery B, Tohyama K, Richardson WD. 2014. Motor skill learning requires active central myelination. Science (80-) 346:318–322.

Micheva KD, Wolman D, Mensh BD, Pax E, Buchanan J, Smith SJ, Bock DD. 2016. A large fraction of neocortical myelin ensheathes axons of local inhibitory neurons. Elife 5:1–29.

Ng MC, Yang YL, Lu KT. 2013. Behavioral and Synaptic Circuit Features in a Zebrafish Model of Fragile X Syndrome. PLoS One 8:1–8.

Al Oustah A, Danesin C, Khouri-Farah N, Farreny MA, Escalas N, Cochard P, Glise B, Soula C. 2014. Dynamics of Sonic hedgehog signaling in the ventral spinal cord are controlled by intrinsic changes in source cells requiring Sulfatase 1. Dev 141:1392–1403.

Pacey LKK, Xuan ICY, Guan S, Sussman D, Henkelman RM, Chen Y, Thomsen C, Hampson DR. 2013. Delayed myelination in a mouse model of fragile X syndrome. Hum Mol Genet 22:3920–3930.

Paluszkiewicz SM, Olmos-Serrano JL, Corbin JG, Huntsman MM. 2011. Impaired inhibitory control of cortical synchronization in fragile X syndrome. J Neurophysiol 106:2264–2272.

Pan L, Zhang YQ, Woodruff E, Broadie K. 2004. The Drosophila fragile X gene negatively regulates neuronal elaboration and synaptic differentiation. Curr Biol 14:1863–1870.

Pan S, Mayoral SR, Choi HS, Chan JR, Kheirbek MA. 2020. Preservation of a remote fear memory requires new myelin formation. Nat Neurosci 23:487–499.

Park H-C, Boyce J, Shin J, Appel B. 2005. Oligodendrocyte specification in zebrafish requires notch-regulated cyclin-dependent kinase inhibitor function. J Neurosci 25:6836–44.

Park HC, Mehta A, Richardson JS, Appel B. 2002. Olig2 Is Required for Zebrafish Primary Motor Neuron and Oligodendrocyte Development. Dev Biol 248:356–368.

Perlman K, Couturier CP, Yaqubi M, Tanti A, Cui QL, Pernin F, Stratton JA, Ragoussis J, Healy L, Petrecca K, Dudley R, Srour M, Hall JA, Kennedy TE, Mechawar N, Antel JP. 2020. Developmental trajectory of oligodendrocyte progenitor cells in the human brain revealed by single cell RNA sequencing. Glia 68:1291–1303.

Pilaz L-J, Lennox AL, Rouanet JP, Silver Correspondence DL, Silver DL. 2016. Dynamic mRNA Transport and Local Translation in Radial Glial Progenitors of the Developing Brain. Curr Biol 26:1–10.

Ravanelli AM, Kearns CA, Powers RK, Wang Y, Hines JH, Donaldson MJ, Appel B. 2018. Sequential specification of oligodendrocyte lineage cells by distinct levels of Hedgehog and Notch signaling. Dev Biol 444:93–106.

Sakry D, Neitz A, Singh J, Frischknecht R, Marongiu D, Binamé F, Perera SS, Endres K, Lutz B, Radyushkin K, Trotter J, Mittmann T. 2014. Oligodendrocyte Precursor Cells Modulate the Neuronal Network by Activity-Dependent Ectodomain Cleavage of Glial NG2. PLoS Biol 12:e1001993.

Sapey-Triomphe LA, Lamberton F, Sonié S, Mattout J, Schmitz C. 2019. Tactile hypersensitivity and GABA concentration in the sensorimotor cortex of adults with autism. Autism Res 12:562–575.

Scott K, O’Rourke R, Gillen A, Appel B. 2020. Prdm8 regulates pMN progenitor specification for motor neuron and oligodendrocyte fates by modulating the Shh signaling response. Development 147:8214.

Shin J, Park HC, Topczewska JM, Madwsley DJ, Appel B. 2003. Neural cell fate analysis in zebrafish using olig2 BAC transgenics. Methods Cell Sci 25:7–14.

Simon C, Götz M, Dimou L. 2011. Progenitors in the adult cerebral cortex: Cell cycle properties and regulation by physiological stimuli and injury. Glia 59:869–881.

Swanson MR, Wolff JJ, Shen MD, Styner M, Estes A, Gerig G, McKinstry RC, Botteron KN, Piven J, Hazlett HC. 2018. Development of white matter circuitry in infants with fragile x syndrome. JAMA Psychiatry 75:505–513.

Tanaka Y, Tozuka Y, Takata T, Shimazu N, Matsumura N, Ohta A, Hisatsune T. 2009. Excitatory GABAergic Activation of Cortical Dividing Glial Cells. Cereb Cortex 19:2181–2195.

Tenzen T, Allen BL, Cole F, Kang JS, Krauss RS, McMahon AP. 2006. The Cell Surface Membrane Proteins Cdo and Boc Are Components and Targets of the Hedgehog Signaling Pathway and Feedback Network in Mice. Dev Cell 10:647–656.

Tervonen TA, Louhivuori V, Sun X, Hokkanen M-E, Kratochwil CF, Zebryk P, Castrén E, Castrén ML. 2009. Aberrant differentiation of glutamatergic cells in neocortex of mouse model for fragile X syndrome. Neurobiol Dis 33:250–259.

Todd PK, Mack KJ, Malter JS. 2003. The fragile X mental retardation protein is required for type-I metabotropic glutamate receptor-dependent translation of PSD-95. Proc Natl Acad Sci 100:14374–14378.

Verkerk AJ, Pieretti M, Sutcliffe JS, Fu YH, Kuhl DP, Pizzuti A, Reiner O, Richards S, Victoria MF, Zhang FP, et al. 1991. Identification of a gene (FMR-1) containing a CGG repeat coincident with a breakpoint cluster region exhibiting length variation in fragile X syndrome. Cell 65:905–914.

Wang F, Ren SY, Chen JF, Liu K, Li RX, Li ZF, Hu B, Niu JQ, Xiao L, Chan JR, Mei F. 2020. Myelin degeneration and diminished myelin renewal contribute to age-related deficits in memory. Nat Neurosci 23:481–486.

Xu H, Dzhashiashvili Y, Shah A, Kunjamma RB, Weng Y lan, Elbaz B, Fei Q, Jones JS, Li YI, Zhuang X, Ming G li, He C, Popko B. 2020. m6A mRNA Methylation Is Essential for Oligodendrocyte Maturation and CNS Myelination. Neuron 105:293–309.e5.

Zalfa F, Eleuteri B, Dickson KS, Mercaldo V, De Rubeis S, Di Penta A, Tabolacci E, Chiurazzi P, Neri G, Grant SGN, Bagni C. 2007. A new function for the fragile X mental retardation protein in regulation of PSD-95 mRNA stability. Nat Neurosci 10:578–587.

